# Comparing kinetic proofreading and kinetic segregation for T cell receptor activation

**DOI:** 10.1101/2023.06.11.544497

**Authors:** Alexander S. Moffett, Kristina A. Ganzinger, Andrew W. Eckford

## Abstract

The T cell receptor (TCR) is a key component of the adaptive immune system, recognizing foreign antigens and triggering an immune response. Competing models exist to explain the high sensitivity and selectivity of the TCR in discriminating ‘self’ from ‘non-self’ antigens, particularly models using kinetic proofreading (KP), kinetic segregation (KS), and combinations of the two. In this paper, we consider the role and importance of KS in TCR activation, using two models: classic KP (cKP), without KS, where antigen-TCR binding is required for activation, and a combination of KP and KS (KS-KP), where only residence within a close contact is required for activation. Building on previous work, our computational model is the first to permit a head-to-head comparison of these models *in silico*. While we find that both models can be used to explain the probability of TCR activation across much of the parameter space, we find biologically important regions in the parameter space where significant differences in performance can be expected. Furthermore, we show that the available experimental evidence may favour the KS-KP model over cKP. Our results may be used to motivate and guide future experiments to determine highly accurate computational models for the TCR.

**Author summary:** The T cell receptor (TCR) is a master of reliable sensing: it detects faint ‘signals’ (rare ligands derived from foreign proteins) over high ‘noise’ (abundant ligands derived from the body’s own proteins) to set T cells on a course to exterminate pathogens and tumours, a process that is central to our immune response. Despite decades of studying TCR signalling, we still do not know how the TCR can be so exceptionally sensitive and accurate. It is widely believed that kinetic proofreading (KP), in which the TCR binds to an antigen and triggers a series of phosphorylation steps prior to activation, plays an important role. However, recent results suggest that kinetic segregation (KS), in which binding is not required, is also important. These models are mutually exclusive, and yet both appear to explain various aspects of T cell activation.

Our work directly addresses this puzzle. We develop a computational modeling framework which can simulate TCR activation by both KP-based and KS-based models, making it possible to compare them *in silico* for the first time. Using this framework, we find conditions under which the two models provide different responses, and we show that the limited experimental evidence to date is consistent with KS, which should motivate further investigation.

## Introduction

In the immune system, the T cell receptor (TCR) is at the heart of T cell function, allowing these cells to discriminate foe from self. TCR signalling is activated when TCRs interact with peptide antigens derived from pathogens or self proteins, for example when presented on peptide-major histo-compatibility complexes (pMHCs) on the membrane of an antigen presenting cell (APC). Activation then triggers a bespoke immune response to the invading pathogen. However, MHCs also present peptides derived from self-proteins, so the TCR has to discriminate rare foreign peptides (as few as 1-10 [1, 2]) from a sea of self-pMHC ligands (10^5^-10^6^ [3]) to allow robust protection from pathogens while minimizing false activation, *i*.*e*. auto-immunity. The TCR therefore operates in a challenging regime, combining high sensitivity with high specificity [4, 5].

The mechanism that enables T cells to process signals so remarkably well has remained elusive – partially because equilibrium thermodynamic processes are insufficient to explain ligand discrimination [6–8]. To address the discrepancy, it was suggested that the TCR uses kinetic proof-reading (KP) [9] to discriminate binding events of different duration [10]. Briefly, KP occurs when ligand binding to a receptor triggers a series of irreversible, energy-consuming biochemical modifications of the receptor until a final, signaling competent state is reached, while intermediate states do not relay the signal. If ligand unbinding results in modifications being removed, this means that only long binding life times result in receptor signaling. Applied to TCR activation through multi-site phosphorylation, KP can account for TCR discrimination; KP models have been found to fit experimental TCR activation data better than models without proofreading steps [5, 11, 12].

The kinetic segregation (KS) model proposes that TCR activation is determined by the TCR’s residence time in so-called close contacts between T cells and APCs, when these cells bring their membranes into close proximity to allow for TCR-pMHC binding [13–15]. Due to the tight inter-membrane spacing, these close contacts are depleted of deactivating phosphatases with large extracellular domains, such as CD45, but retain the TCR kinase Lck needed for TCR phosphrylation and activation [16–18]. TCRs will activate if they remain in a close contact for a sufficiently long time so that they are fully phosphorylated, with ligand binding only serving to enhance residence times by preventing TCRs from diffusing out of the close contact. Among other work supporting the KS model, TCR elongation was shown to prevent TCR signaling [19], while TCRs were observed to activate ligand-independently within sufficiently large close contacts with a decreased CD45 to Lck ratio [18].

Although KP and KS are widely different, there is little agreement in the literature over which model, or combination of models, best explains TCR function. There is particular uncertainty over the role and importance of KS, despite experimental evidence [13, 20]. Compounding the problem are two issues. First, the models are seemingly incompatible: classic formulations of KP assume that the TCR is reset to its initial state by ligand unbinding, while KS assumes that phosphorylation can occur as long as the TCR is in the close contact, whether ligand bound or not, and that resetting only occurs when a TCR leaves the close contact. Second, KP and KS models are not generally formulated in a way that can be directly compared: for example, KS requires careful handling of diffusion within the close contact (see e.g. [21]), while receptor diffusion is ignored in classic KP models.

Here, we address these issues by developing a framework for directly comparing KP and KS models, and by using this framework to test how performance differs for these models and their combinations. To do so, we first develop a reformulation of KP that is compatible with the KS model, and explore how well this new model can account for the TCR’s remarkable ligand discrimination (KS-KP model). Second, to compare the KS-KP model to a more classic KP model (cKP model), we have developed a new computational framework for describing TCR activation, capable of describing a classic receptor activation-KP model in context of the close contacts postulated by the KS model. We find that both models predict similar TCR sensitivities. However, we also find that the KS-KP model predicts a much higher probability of activation than the cKP model in the absence of non-self ligands, for realistic TCR phosphorylation rates. While our modelling suggests that both KS-KP and cKP predict similar patterns of TCR activation across a wide range of parameters, important differences are predicted in some ranges of biologically important situations, such as when small numbers of non-self antigens are present. We also find that some simplified assumptions used in earlier models may further obscure the difference between the models. We conclude that the role of KS in general, and the cKP and KS-KP models in particular, may heretofore have been difficult to distinguish through the mere comparison of downstream metrics of T cell activation in the absence of biochemical data. Further, we show how recent biochemical experiments may support KS-KP, and that our model may indicate where best to look in order to distinguish between cKP and KS-KP.

## Results

### Computational modeling of KS-KP and cKP

A key contribution of our work is a mathematical framework and computational model that allows a direct comparison between the classic KP (cKP) model, and a kinetic segregation-kinetic proofreading (KS-KP) model (Fig 1). In our activation model, described in detail in the Materials and Methods section, we explicitly consider the close contacts formed between T cells and APCs. We allow individual TCRs to diffuse into and out of the close contact by Brownian motion, binding to and unbinding from both ‘self’ and ‘non-self’ ligands (i.e. agonists) while being in the close contact. We assume that ligand-bound TCRs are unable to leave the close contact, since we define the close contact as the region in which the cell membranes of the two contacting cells are in sufficiently close apposition to allow a pMHC-TCR complex to form across the two contacting cells. Each TCR complex has *n* phosphorylation sites, and we consider a T cell to be activated once all *n* phosphorylation sites on *at least one* TCR complex are phosphorylated.

**Fig 1.**
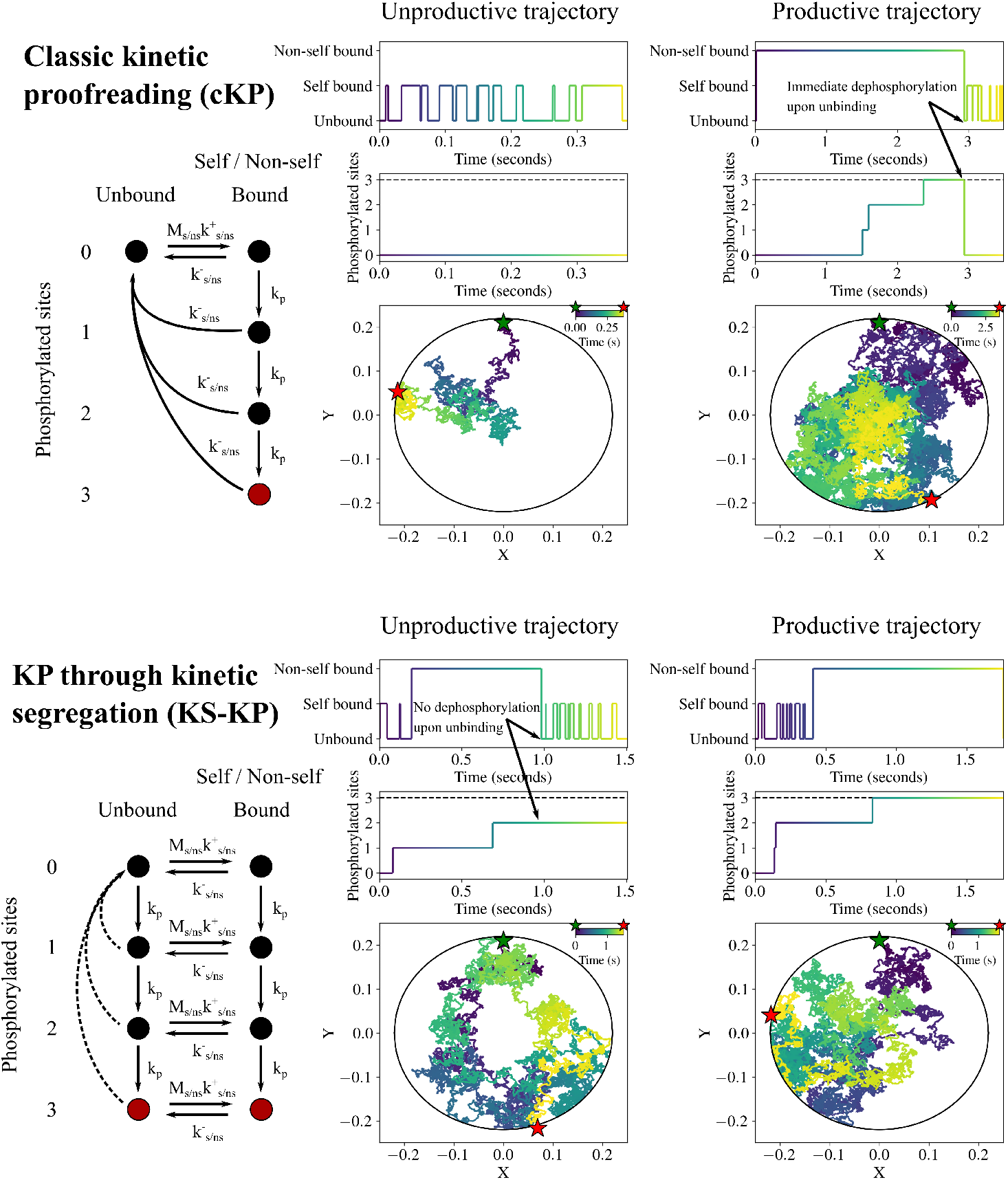
An overview of our computational models of T cell receptor (TCR) activation in the close contact context. The classic kinetic proofreading (cKP) model is shown in the top half of the figure, depicted with three phosphorylation sites (*n* = 3) in the example. On the left is a simplified diagram of the cKP model, where a TCR can be phosphorylated when bound to self or non-self ligands and is rapidly dephosphorylated upon ligand unbinding. The active states, where all three phosphorylation sites are modified, are shown in red. An unproductive trajectory (where the TCR does not reach the active state before leaving the close contact) is shown in the middle column while a productive trajectory is shown in the right column. Note that in the unproductive trajectory, the TCR only binds to self ligands, with a rapid off rate preventing phosphorylation. In the productive trajectory, a long period of time where the TCR is non-self bound leads to all three sites being phosphorylated while the TCR is immediately fully dephosphorylated upon ligand binding. The kinetic segregation (KS-KP) model is shown in the bottom half of the figure. The diagram on the left shows how with the KS-KP model, phosphorylation can take place whether a ligand is bound or not and can only be reversed when the TCR leaves the close contact (depicted with dashed arrows). Accordingly, in both the unproductive and productive trajectories shown for KS-KP, phosphorylation events occur for both bound and unbound TCRs. KS-KP is still sensitive to ligand concentrations, as ligand binding influences the time that a TCR spends in the close contact before leaving.

Unlike earlier related models [21, 22], we apply different (de)phosphorylation rates depending on whether the ligand is bound or unbound. In cKP, the unbound phosphorylation rate is zero, and the unbound dephosphorylation rate is effectively infinite, resulting in an immediate reset. Meanwhile, in KS-KP, the (de)phosphorylation rate is independent of the binding state in the close contact, and the time scale for reset depends on binding only insofar as binding increases the TCR’s residence time in the close contact, as effective dephosphorylation only occurs once a TCR has diffused out of the close contact. (As a check of the accuracy of our diffusive model, in the SI we show that the residence time in the close contact *τ* is dependent on non-self off rate, indicating that the binding dynamics of the ligand modify its diffusion, as expected; see Fig S1). The allowance for this key difference in our computational framework allows us to directly compare cKP with KS-KP; it also admits a spectrum of models in between the two extremes, a subject for future work.

As we model only the very first steps of TCR activation, we assume that TCRs do not interact. While formation of TCR clusters is known to play an important role in T cell activation [23, 24], experimental data also suggests that a single TCR is sufficient to activate a T cell [1, 2], consistent with our assumption that the T cell is activated once all sites on a TCR are phosphorylated.

Finally, as a check for consistency of our framework with previous models, we obtain from our simulations the probability of a single TCR remaining in the close contact as a function of time; we then compare it with the same quantity calculated from a partial differential equation (PDE) model adapted from Fernandes *et al*. [22]. The simulation results match reasonably well with those from the PDE model (Fig S2). We may therefore assume that the results we obtain are comparable with results obtained from earlier related models.

### Subtle differences in number of phosphorylation sites result in highly similar T cell activation being predicted by both cKP and KS-KP models

We first investigated how the number of proofreading steps, *n*, affect the activation probability for different non-self/self ligand ratios in both KS-KP and cKP models at a fixed phosphorylation rate (*k*_*p*_ = 1 *s*^*−*1^), using values for *n* in the range of 2-4 (Fig 2).

**Fig 2.**
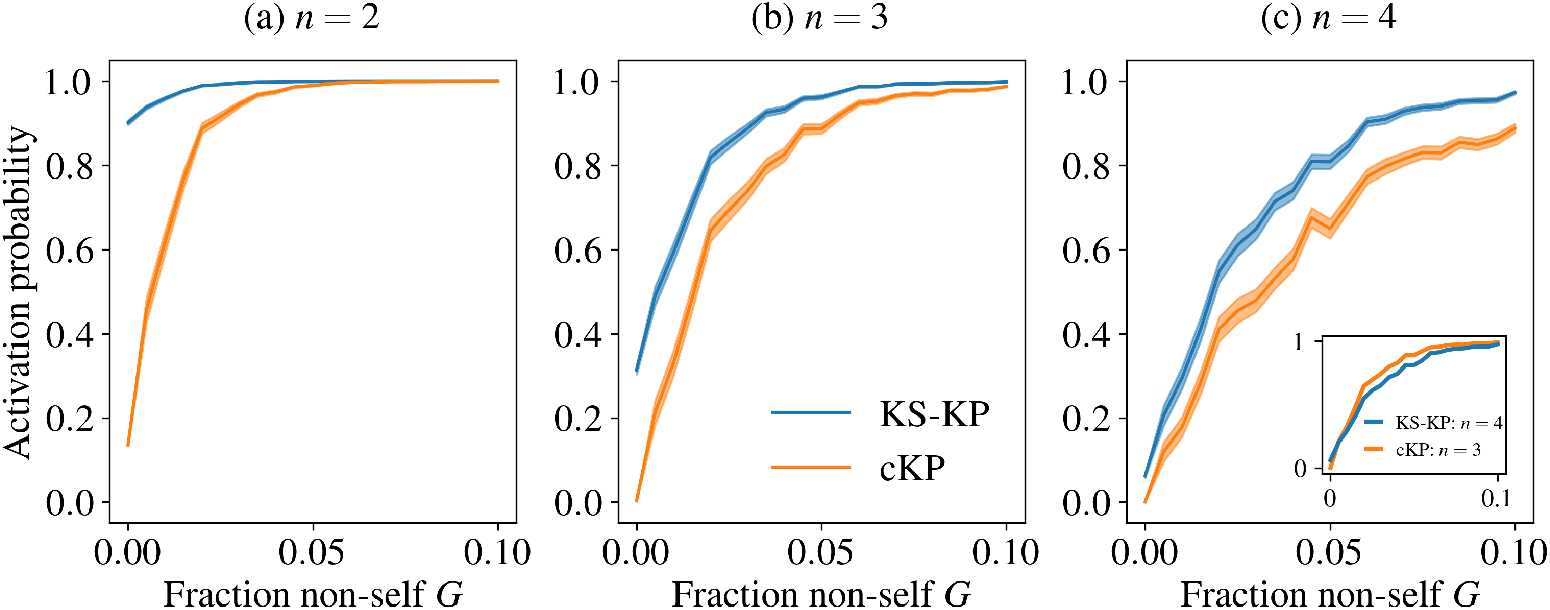
Activation probability as predicted by KS-KP is much more strongly dependent on number of phosphorylation steps *n* than cKP. Activation probability is plotted against the fraction of non-self ligand for *n* = 2 (a), *n* = 3 (b), and *n* = 2 (c). As the number of steps increases, activation probability decreases, as it is less likely for the TCR to remain in in the close contact (in KS-KP) or bound to the ligand (in cKP) until complete phosphorylation. Phosphorylation rate is set to *k*_*p*_ = 1 s^*−*1^; examples for other values of *k*_*p*_ are found in SI, Fig S4. *Inset in (c):* KS-KP and cKP yield similar results when compared at slightly different values of *n*. The KS-KP *n* = 4 line is superimposed on the cKP *n* = 3 line; the responses are similar, suggesting that solutions for both KS-KP and cKP can be readily found. Further examples are provided in SI, Fig S3. Error bars are standard deviations, calculated as described in the Materials and Methods. See Table S1 for the parameters used in this figure.

We note that while proofreading steps may be directly linked to phosphorylation sites on the TCR, we use *n* in the range that was previously found to well explain T cell activation data [5] as opposed to the full number phosphorylation sites on a TCR (i.e. 10). We found that generally, the activation probability predicted by KS-KP was much more strongly dependent on *n* as that predicted by the cKP model. Interestingly, at *n* = 3, the activation probability for KS-KP and cKP models appeared to diverge less from each other than for other values of *n* explored. Strikingly, KS-KP and cKP yielded very similar results when compared at slightly different values of *n* (inset to the *n* = 4 subfigure in Fig 2, overlaying the KS-KP *n* = 4 and cKP *n* = 3 lines at *k*_*p*_ = 1 *s*^*−*1^). In addition to this example, similarly coincidental curves are readily found if both models are compared for different combinations of *n* and *k*_*p*_, which we report in the SI (Fig S3). These results suggest that KS-KP and cKP may be difficult to distinguish through downstream metrics of T cell activation as e.g. in [5] without careful consideration of the biochemical details.

In the SI, we give results where the phosphorylation rate *k*_*p*_ is varied. (Fig S4). We found that for a lower phosphorylation rate (*k*_*p*_ = 0.1 *s*^*−*1^), KS-KP performed slightly better than cKP in terms of sensitivity and discrimination. Meanwhile, at a higher phosphorylation rate (*k*_*p*_ = 10 *s*^*−*1^), KS-KP was entirely unable to distinguish between different non-self fractions for all *n* tested, while cKP was only able to do so well at n = 4.

### For realistic activation time distributions, KS-KP results in higher activation probabilities in the absence of non-self ligands than cKP

Here we consider activation probability with respect to fraction non-self *G* (Fig 3) and with respect to close contact lifetime *t*_*c*_ (Fig 4), comparing KS-KP and cKP along with two different models for activation, described as follows. First, assuming first-order kinetics, TCR phosphorylation times (for each step) are exponentially distributed. If each of the *n* steps occurs at a similar rate, it is reasonable to approximate the total time (i.e., the time until full phosphorylation or activation time) as an Erlang-*n* distribution (see Materials and Methods; Eq. 23). Second, earlier work by Fernandes *et al*. [22] used a simplified model for TCR activation, in which any TCR that remained in the close contact for at least 2 s was immediately and fully activated, i.e., the effective probability density function for the activation time has a Dirac delta function at *t* = 2 s. We thus refer to this earlier model as the Delta model.

**Fig 3.**
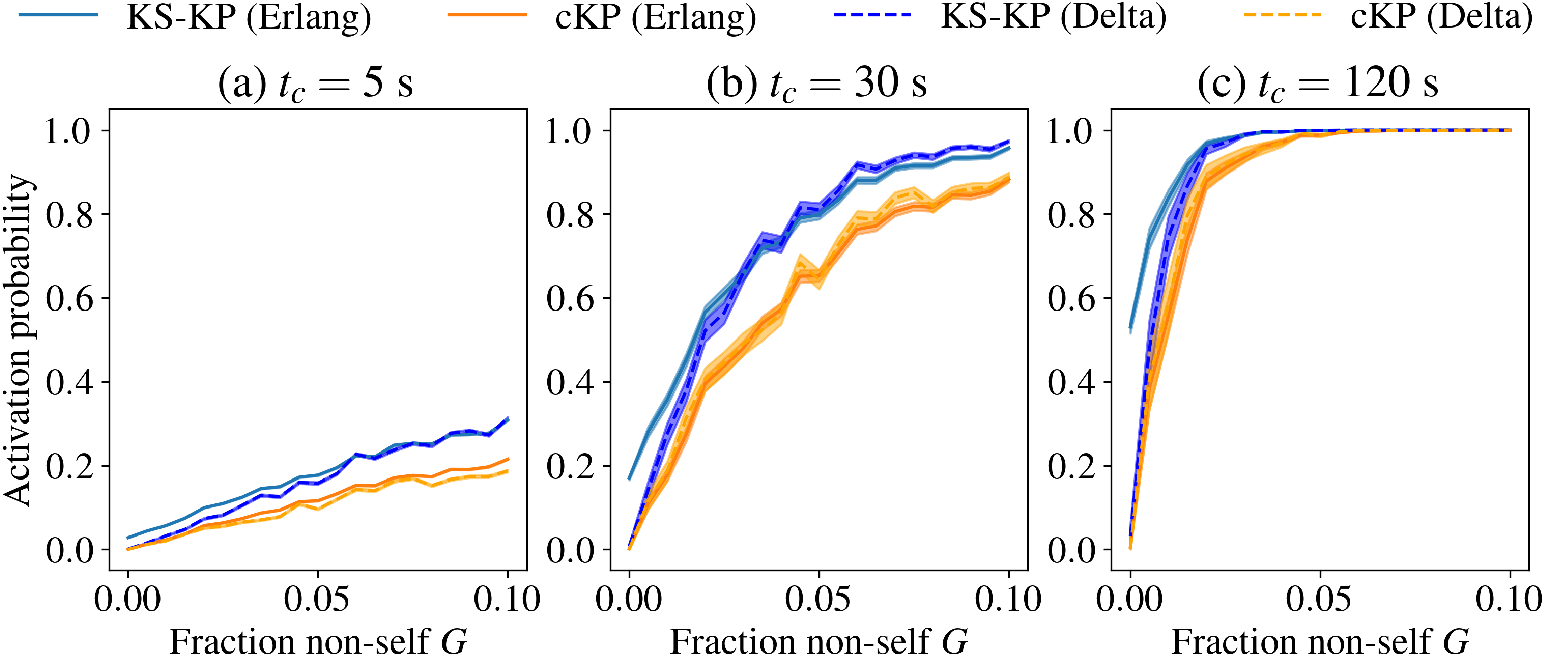
KS-KP leads to increased activation probabilities as compared to cKP, especially at low non-self fractions. Subfigures (a-c) show the overall activation probability *P*_*a*_(*t*_*c*_) as a function of the fraction of non-self ligands for varying close contact lifetimes, *t*_*c*_. For the Erlang distribution model of phosphorylation, KS-KP drastically increases the activation likelihood over cKP when the non-self ligand fraction is very low, suggesting that KS-KP is not able to discriminate between self and non-self ligands as well as cKP when non-self ligands are few. The phosphorylation scheme in which TCRs are instantly, fully phosphorylated at 2s (labeled as Delta, for the distribution *δ*(*t −* 2s)) leads to very similar activation probabilities for the cKP scheme, as compared with an Erlang distribution (Eq. 23) with *n* = 3 and *k*_*p*_ = 1*s*^*−*1^. For the KS-KP scheme, however, the Erlang phosphorylation model predicts drastically higher activation than the Delta model at low non-self ligand fractions. Examples for additional values of *t*_*c*_ are given in the SI, Fig S5. Error bars are standard deviations, calculated as described in the Materials and Methods. See Table S1 for parameters used in this figure.

**Fig 4.**
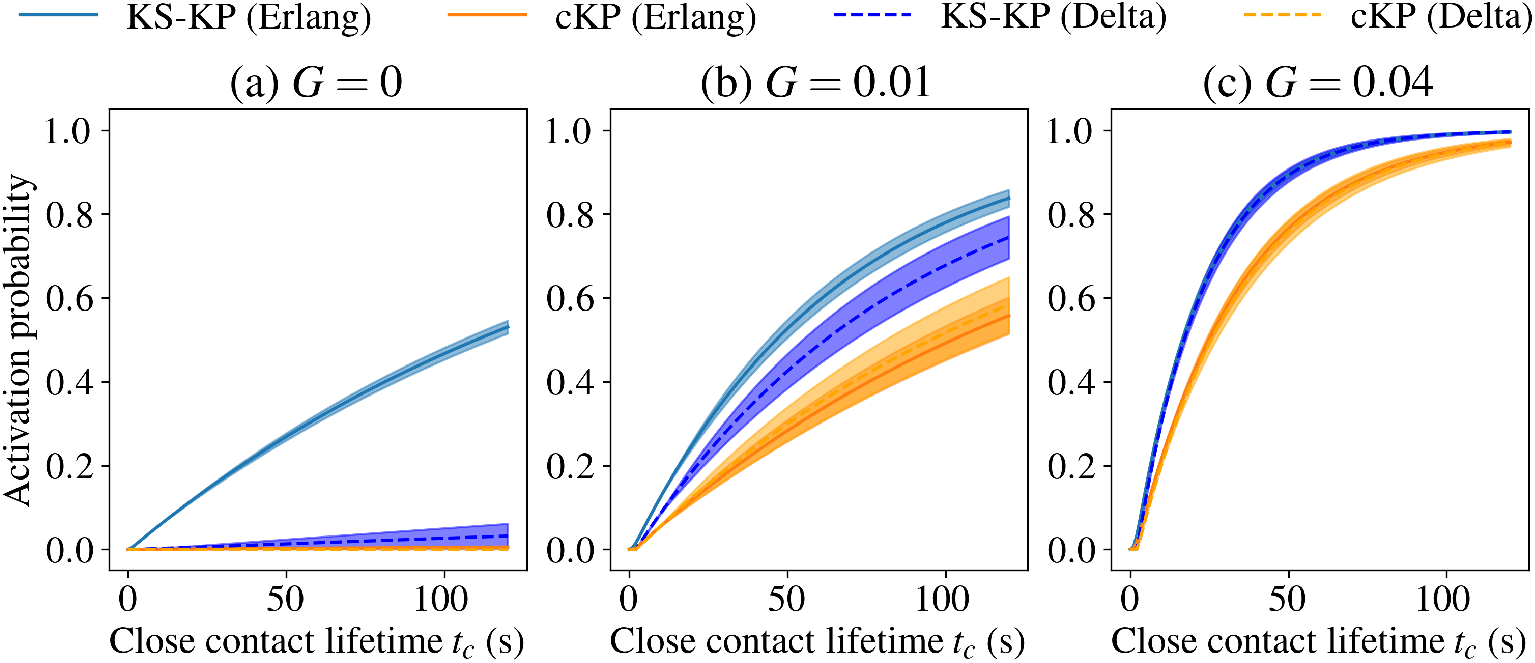
At low non-self ligand fractions *G*, the Erlang distribution model (Eq. 23) with KS-KP differs from KS-KP with the Delta model, as well as cKP under either model (cf. Fig 3). (a-c) Overall activation probability *P*_*a*_(*t*_*c*_) as a function of close contact lifetime *t*_*c*_ for different fractions of non-self ligands, *G*. The difference between assuming a Delta or Erlang distribution is most pronounced at *G* = 0 (a), and rapidly shrinks with increasing *G* ((b), (c)). At *G* = 0, only the KS-KP Erlang model shows significant probability of activation. Examples for additional values of *G* are given in the SI, Fig S6. Error bars are standard deviations, calculated as described in the Materials and Methods. See Table S1 for parameters used in this figure.

In our KS-KP model, when we compared the predicted T cell activation for the more realistic Erlang distribution to the Delta model used in [22], we found that the Delta model leads to a marked underestimation of KS-KP T cell activation at low non-self ligand fractions *G* (Fig 3, Fig 4 (a-b), Fig S5 (a-d)). Importantly, at these low values of *G*, the performance of KS-KP with the delta activation model is similar to the performance of cKP in combination with either activation model. At high values of *G*, there is little difference in performance, and all models indicate a high probability of activation for reasonably large *t*_*c*_ (Fig 3, Fig 4 (c), Fig S5 (e-i)).

A key point is that the activation probability of the KS-KP Erlang model at low *G* is consistent with experimentally observed behaviour of the TCR, where activation in the absence of ligand was observed [18, 19]. This is especially clear in Fig 4 (a), where the KS-KP Erlang model has a significant activation probability; the other models in the figure show zero activation probability. Our model predicts that this difference should rapidly diminish as *G* increases: In Fig 4 (b), at *G* = 0.01, all four models show significant probability of activation, while in Fig 4 (c), the gap between the Erlang and Delta models diminishes to practically nil at *G* = 0.04. Still, the case where *G→* 0 is biologically important, and none of the competing models (i.e. KS-KP Delta, cKP Delta, or cKP Erlang) are consistent with activation in the absence of non-self ligands.

Moreover, as *G →*0, it is interesting to note that the difference between cKP and KS-KP is minimal unless the Erlang model is used (Fig 4 (a)); that is, using the Delta model obscures the difference between these models at small *G*. In the KS-KP model, the difference between the Erlang and Delta models at low *G* values can be rationalized by the fact that a single TCR is highly unlikely to remain within the close contact for 2 s or longer when unbound (as is clear in Figs S1 and S2). Thus, with instant, full phosphorylation only at 2 s, a free TCR is very unlikely to activate. In contrast, an Erlang distribution with *n* = 3 and *k*_*p*_ = 1 *s*^*−*1^ has sizable density below 2 s, meaning full phosphorylation of free TCRs with residence times of less than 2 s is possible, resulting in more ligand-independent activation events. Another corollary of the sizable density of phosphorylated TCRs below 2 s of the Erlang model is that Erlang model gives positive activation probability for any *t*_*c*_ *>* 0 while the Delta model implies an activation probability of zero for *t*_*c*_ *<* 2 s; an illustration of this difference is provided in the SI (Fig S7).

### KS-KP and cKP predict distinct responses to different non-self ligand off rates

Finally, in Figs 5 and 6, we examine the response of the KS-KP and cKP models to non-self ligands with different off rates. Non-self ligand off rates have been shown to correlate with TCR activation [8, 25, 26], a fact that has been used as evidence of KP. In this series of results, we test non-self ligand off rates of 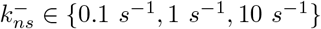 and we use a self ligand off rate of 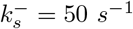 for all simulations; that is, the weakest binding non-self ligand we examine 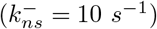 has a 5-fold smaller dissociation constant than the self ligand [22].

**Fig 5.**
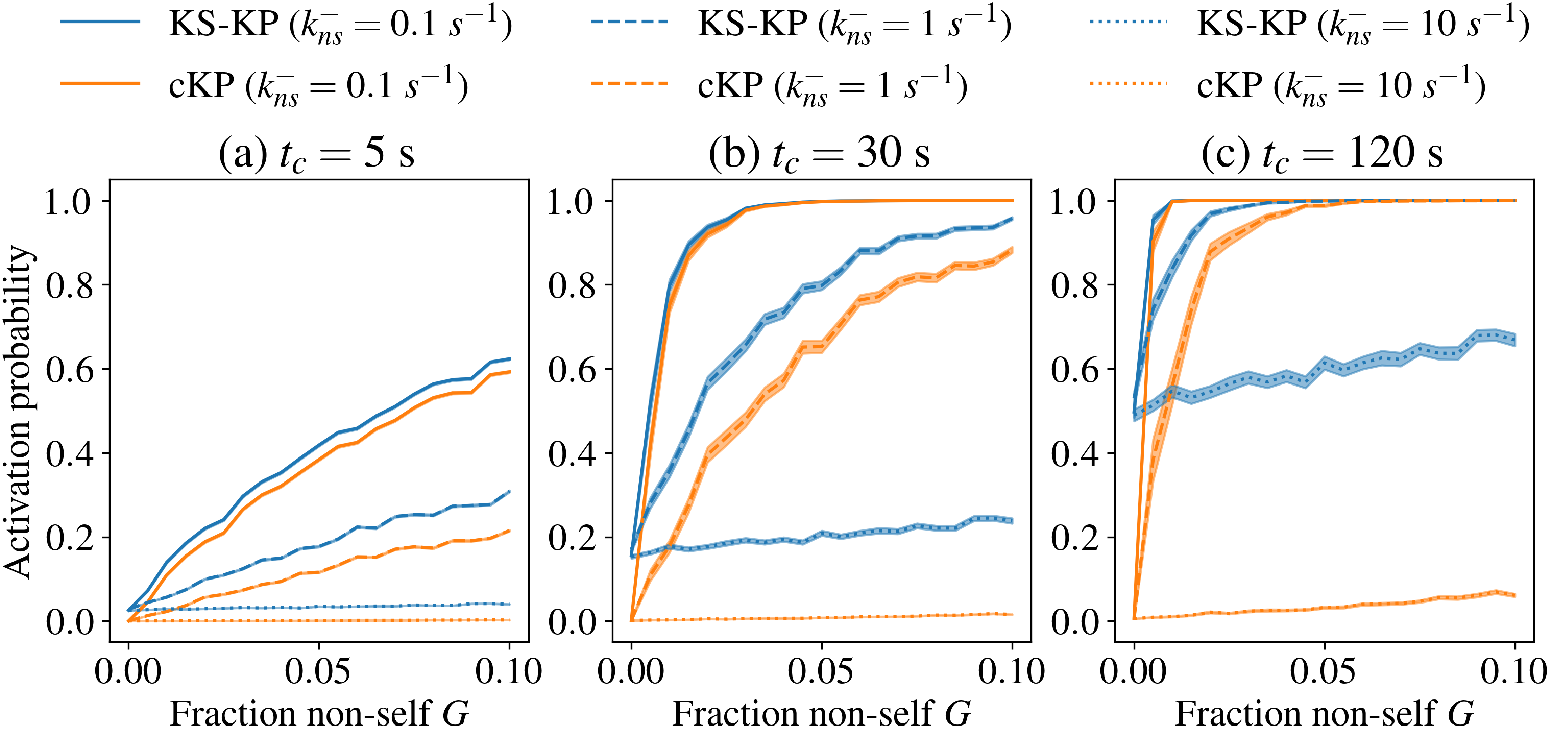
cKP is able to mount more distinct responses than KS-KP to non-self ligands with different off rates, especially at higher close contact lifetimes. (a)-(c) Activation probability for TCRs with the KS-KP and cKP models with 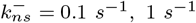, and 10 *s*^*−*1^ as a function of non-self fraction *G* at the indicated close contact lifetime. Examples for additional values of *t*_*c*_ are provided in the SI, Fig S8. Error bars are standard deviations, calculated as described in the Materials and Methods. See Table S1 for parameters used in this figure.

**Fig 6.**
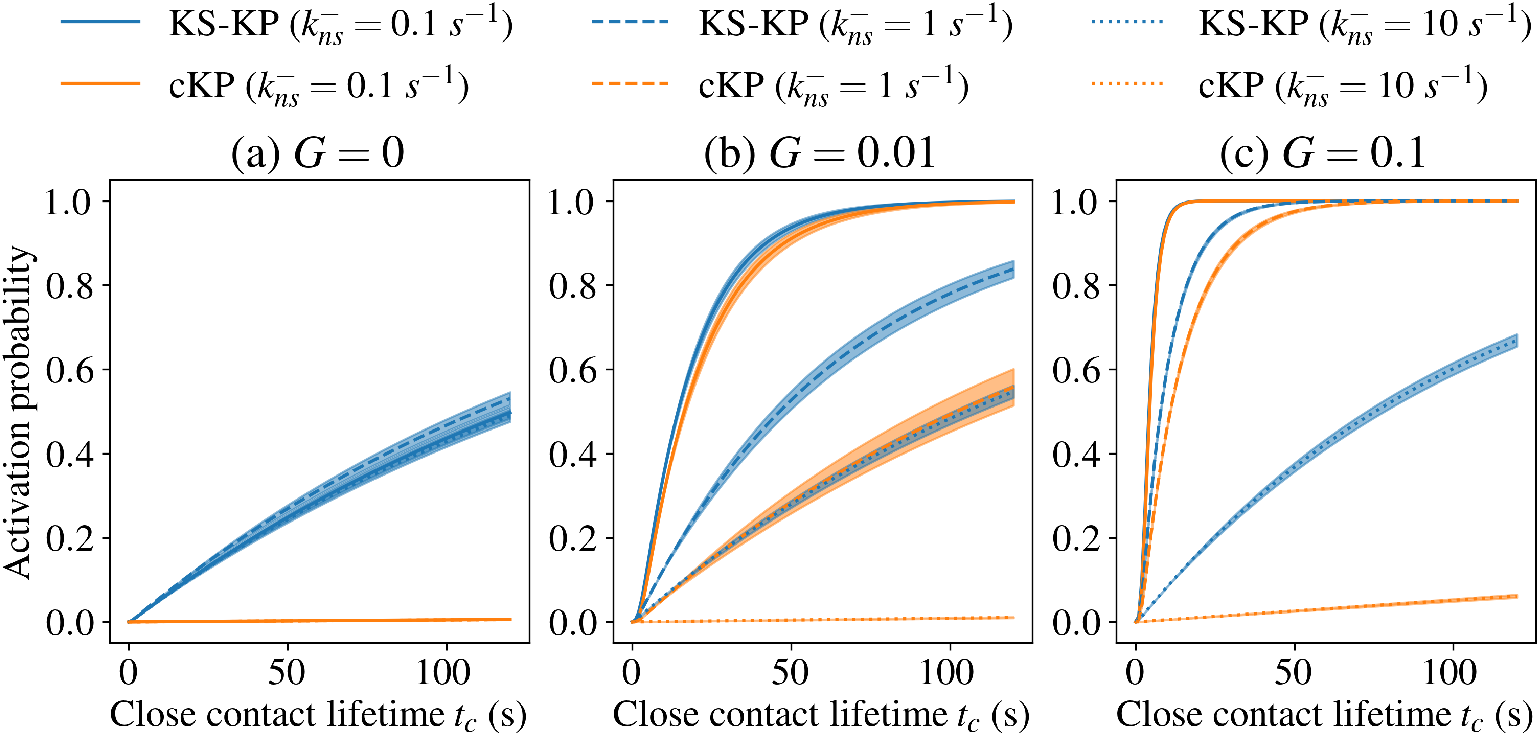
Activation probability for TCRs with the KS-KP and cKP models with 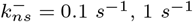 and 10 *s*^*−*1^ as a function of close contract lifetime at the indicated non-self fraction *G*. At low non-self fraction *G* ((a), (b)), the activation probability for KS-KP (blue lines) at all values of 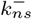 is significantly greater than that for cKP (orange lines). At high non-self fraction (c), a strong difference only exists for large 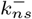 cf. Fig 5. Examples for additional values of *G* are provided in the SI, Fig S9. Error bars are standard deviations, calculated as described in the Materials and Methods. See Table S1 for parameters used in this figure.

We first consider activation probability versus close contact lifetime *t*_*c*_. For small values of *t*_*c*_.activation probability varies with 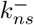 (as expected), but the difference between KS-KP and cKP is slight (Fig 5 (a), *t*_*c*_ = 5 s). At larger values of *t*_*c*_ (Fig 5 (b,c), *t*_*c*_ = 30, 120 s), there are significant differences between KS-KP and cKP at smaller 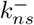. We subsequently consider activation probability versus non-self fraction *G*. Similar to the previous section, large differences between the models are also observed at small *G*, which are more clearly illustrated in Fig 6. At non-self fraction *G* = 0, only KS-KP gives a nonzero activation probability (Fig 6 (a)), and for a small value of *G* (e.g., *G* = 0.01), there remain significant differences between KS-KP and cKP (Fig 6 (b)) (cf. Fig 4 (b-c)). For larger *G* (e.g., *G* = 0.1), the differences are reduced at smaller values of 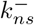 but they remain at higher values of 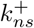 (Fig 6 (c)), consistent with the results in Fig 5. In all figures, when 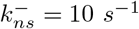 (corresponding to a very weakly binding ligand), a large difference is observed between KS-KP and cKP across all non-self fractions *G*, except at the smallest values of *t*_*c*_.

In summary, we find similar behavior between the two models as we change the non-self ligand off rate 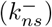, except that (as before) the KS-KP model predicts notable, close contact lifetime-dependent minimum activation probabilities achieved at zero non-self ligand fractions (Fig 6). This is similar to the behaviour observed in Fig 4, and is again consistent with experimental observations of activation in the absence of non-self ligands [18, 19]. While KS-KP displays a close contact lifetime-dependent increase in minimum activation probability, the activation probability for cKP increases very little.

### KS-KP model better distinguishes between experimentally characterized self and non-self ligands

In order to compare our theoretical results for the cKP and KS-KP models to experimental data, we used published EC_40_ and EC_50_ values for seven ligands with known on and off TCR binding rates [25, 26] (see Table S2). In aggregate, these experiments provide information about the *overall* T cell activation probability for three self ligands and four non-self ligands each in the absence of any other ligand. In order to compare the cKP and KS-KP models to these results, we ran simulations and calculated activation probabilities as before, using the corresponding experimental on and off binding rates for each ligand. For self ligands we used *G* = 0 and for non-self ligands we used *G* = 1, while varying the overall ligand concentration, *M*. We estimated EC_40_ and EC_50_ values (according to the quantity available for each ligand, from here on referred to as EC_40*/*50_) by finding the value of *M* for which the activation probability was 40% or 50% maximal, respectively.

In Fig 7, we show the results of our estimated EC_40*/*50_ values compared with the corresponding experimental values for both the cKP and KS-KP models. We emphasize that the small number of data points makes it difficult to draw conclusions. While we do not expect our proxy for T cell activation (*P*_*a*_) to quantitatively predict experimental EC_40*/*50_ values, both the cKP and KS-KP models were able to distinguish self from non-self ligands, with lower EC_40*/*50_ values for all non-self ligands.

**Fig 7.**
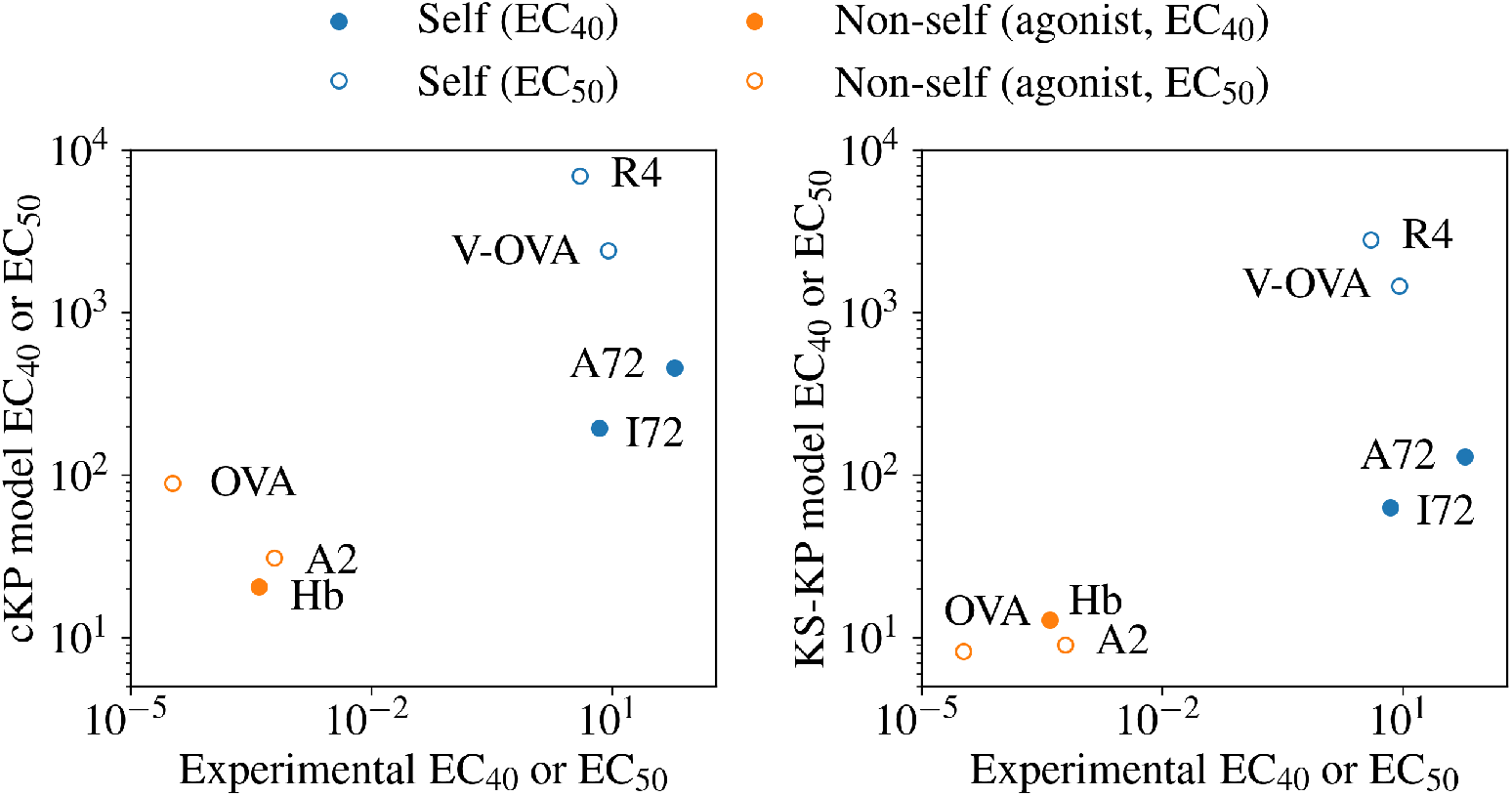
Comparison of experimental EC_40_ and EC_50_ values for seven ligands with theoretical predictions from the cKP and KS-KP models. The model EC_40_ and EC_50_ values are the ligand concentrations at which activation probability is equal to 0.4 and 0.5, respectively, with *G* = 0 for self ligands and *G* = 1 for non-self ligands to match experimental conditions where a single ligand type is present. Both the cKP and KS-KP models differentiate self from non-self ligands, while the KS-KP half-maximal activation probability ligand concentrations are lower than the cKP predictions, consistent with the results in previous sections. The seven ligands considered, along with the corresponding binding and unbinding rates, are listed in Table S2. See Table S1 for the parameters used in this figure.

## Discussion

In this work, we have developed two reformulations of KP models for TCR signaling: one that is fully compatible with the KS model of TCR signaling (KS-KP model) and a more classic KP model in the context of the close contacts postulated by the KS model (cKP model). Considering existing models in the literature, our KS-KP model is generalized so that phosphorylation rates while bound can be different from those while unbound (compared with [21]), and has a more accurate model of activation time (compared with [22]), while applying a diffusion framework to cKP to allow direct comparison with models including KS (i.e., KS-KP). By directly comparing cKP and KS-KP models of TCR activation, we have found that both models of early steps in TCR signaling predict similar TCR discrimination and sensitivity. However, a key difference between the models is that KS-KP results in a close contact lifetime-dependent minimum activation probability, even in the absence of non-self ligands (agonists).

Over a wide parameter range, cKP and KS-KP models make similar predictions. This is likely due to the fact that the kinetic segregation mechanism in KS-KP also implements a form of kinetic proofreading, where a reset to the fully dephosphorylated state occurs upon the TCR complex diffusing out of the close contact (instead of upon ligand dissociation, as in cKP). Our modelling of cKP and KS-KP in a close-contact zone context suggest that all else being equal, KS-KP leads to higher activation probabilities than cKP along with sizable, non-zero activation probabilities without non-self ligands present. This positive minimal activation probability results from any non-self ligand increasing the activation probability as long as 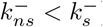) and results in worse differentiation between non-self ligands (agonists) with varied off rates in the KS-KP than cKP model. However, we note that for lower phosphorylation rates (*k*_*p*_ = 0.1 *s*^*−*1^) KS-KP performed better than cKP in both terms of sensitivity and discrimination as for this rate, full phosphorylation of the TCR (and hence activation) cannot be reached before ligand dissociation in cKP. Conversely, at fast phosphorylation rates (*k*_*p*_ = 10 *s*^*−*1^), KS-KP was entirely unable to distinguish between different non-self fractions for all *n* tested as at this rate all TCRs are fully phosphorylated before they leave close-contacts, even for residence times expected for free TCRs. For other parameters, KS-KP and cKP are generally similar at low *t*_*c*_, low-to-moderate 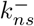, or moderate-to-high *G*, while significant differences are seen at high 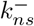 and low *G*, with these differences being largely due to the KS-KP model’s phosphorylation in the close contact region independent of ligand binding. Experiments designed to distinguish the two models should therefore be in this parameter regime.

In this light, contemporary experimental evidence observed for different TCR signaling models is hard to interpret unambiguously. For example, recently published experimental work that has been interpreted as evidence for cKP [11, 12] would also be consistent with KS-KP if under the conditions used for these studies phosphatases are excluded from regions of TCR-ligand interactions. Other experimental evidence indicates that TCRs may activate even with no non-self ligands present, [18, 19], which is more likely under KS-KP than cKP, but the high specificity seen in [1, 2] is more readily explained by cKP. An advantage of our computational approach is that both KS-KP and cKP might be viewed as extreme points on a spectrum of possible behaviours: for example if ligand binding changes *k*_*p*_ but is not essential for it, such a model would exist between KS-KP and cKP, and could be evaluated in our framework in future studies.

In conclusion, our results may partly explain the ambiguity surrounding the models for TCR activation, as without direct evidence for the number of phosphorylation steps and the rate of phosphorylation, and without experiments that recreate the close contact context in which TCRs are activated, our results suggest that distinguishing between the KS-KP and cKP models through downstream signaling events may be difficult. Moreover, the differences only emerge as the models increase in fidelity (for example, consider the difference between the Erlang and Delta models of activation at *G* = 0, Fig 4 (a)). On the other hand, our results point to clear differences that carefully designed experiments could measure. For example, if TCR activation and discrimination was measured for different phosphorylation rates, e.g. making use of kinetic mutants of Lck [27, 28], ligand discrimination should be much more affected for cKP than KS-KP. These promising potential directions of future experimentation, combined with consistency between our model and experimental findings on TCR activation, is encouraging to future experimental work. We hope that our results will facilitate further investigation into TCR signalling mechanisms, perhaps leading to a synthesis of two views of T cell activation (KS and cKP) that are often viewed as being mutually exclusive.

## Materials and Methods

### Stochastic simulation of TCR diffusion in a close contact

We wish to directly compare the behavior of a typical kinetic segregation model with that of a kinetic proofreading model in the context of a T cell-APC close contact. In KS models, a single TCR can be phosphorylated as long as it is in the close contact, regardless of whether it is bound to a ligand or not, and dephosphorylation occurs only upon leaving the close contact. In KP models, TCRs can be phosphorylated only when bound to a ligand and are rapidly dephosphorylated upon ligand unbinding. In order to capture the behavior of both KS and KP models, we used stochastic simulations of a TCR in a close contact. We simultaneously modeled Brownian motion of the TCR and its ligand binding dynamics, where interaction between the two processes has significant effects on activation probability for cKP and no effect on KS-KP activation beyond extending close contact lifetimes. We will explain this point further below. Within the close contact, we model TCR motion according to

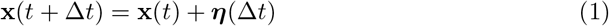

where **x**(*t*) is the 2-dimensional Cartesian coordinates of the TCR at time *t*, ∆*t* is the time step used, and ***η***(∆*t*) is total random motion undergone by the TCR over ∆*t*. The random motion ***η***(∆*t*) is independent between successive time steps and between dimensions, and is normally distributed with

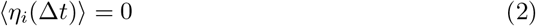

and

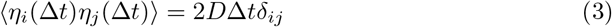

for dimensions *i, j* ∈ {1, 2}, where *D* is the diffusion coefficient and *δ*_*ij*_ is the Kronecker delta function.

Using the same time step ∆*t*, we simulated the binding dynamics according to the transition probability matrix

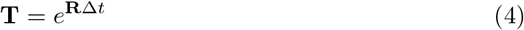

defined with the rate matrix

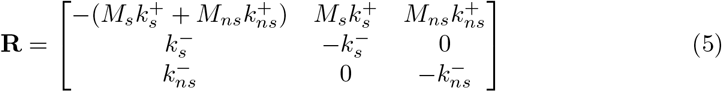

where the first row corresponds to transitions out of the unbound state, the second row to transitions out of the self bound state, and the third row to transitions out of the non-self bound state. The concentrations of self and non-self ligand are written as *M*_*s*_ and *M*_*ns*_, where the overall ligand concentration is set as *M* and *M*_*s*_ and *M*_*ns*_ are determined by the non-self fraction *G*

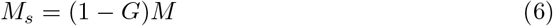

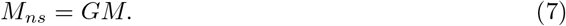

We write the biochemical state of the TCR at time *t* as

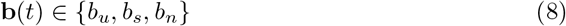

where *b*_*u*_ is the unbound state, *b*_*s*_ is the self bound state, and *b*_*n*_ is the non-self bound state. We modeled the close contact as a 2-dimensional disk Ω with radius *R* = 220 nm centered at the origin,

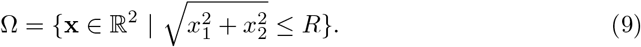

In keeping with previous KS models of TCR dynamics, we assume that the TCR can only leave the close contact when in the unbound state, so that when the TCR was in the *b*_*s*_ or *b*_*n*_ biochemical states, the Brownian motion step was repeated until the TCR remained within Ω at the next time. When in the *b*_*u*_ state, a Brownian motion step that led to the TCR leaving Ω ended the simulation run.

We initialized all simulations with **x**(0) = [0 nm, 210 nm] and **b**(0) = *b*_*u*_, in the unbound state. We ran each simulation until either the TCR left the close contact or until 10 s of model time had elapsed, and repeated each simulation 10,000 times. We performed these simulations using random sampling with the Numpy package in Python [29].

### PDE model of TCR diffusion in a close contact

We independently calculated the probability of a TCR remaining in a close contact after time *t* by adapting a previously published model of TCR diffusion, employing a coupled system of partial differential equations (PDEs) [22].

As with the stochastic simulations, we model the close contact as a disk of fixed radius *R* = 220 nm (as in in Eq. 9), which individual TCRs can diffuse in and out of. First, we consider the diffusion of an individual TCR from the instant it has entered the close contact. Starting at this instant, we are interested in calculating the probability that the TCR then exits the close contact after a given amount of time has passed. We write locations within the close contact as **x** = [*x*_1_, *x*_2_] under the constraint 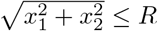. A TCR in the close contact can exist in unbound, self-bound, and non-self-bound form. The two-dimensional diffusion coefficients can in principle differ between unbound (*D*_*T*_) and bound (*D*_*C*_) TCRs, but here are identical. Once a TCR reaches the boundary of the close contact, it is absorbed (leaves the close contact) if unbound to a ligand or is reflected if ligand-bound. We write the boundary as

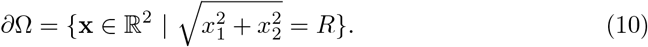

Our goal is to find the probability distributions of unbound, self-bound, and non-self-bound TCRs over Ω as a function of time since TCR entry to the close contact.

We can find these distributions by solving a coupled system of PDEs for the probability density of unbound, *T* (**x**, *t*), self-bound, *C*_*s*_(**x**, *t*), and non-self-bound, *C*_*ns*_(**x**, *t*), TCR at **x** at time *t*

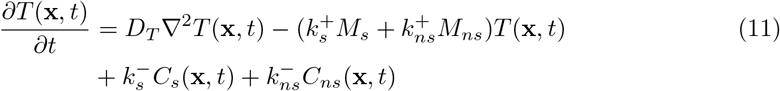

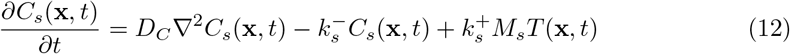

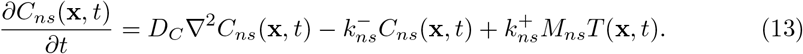

Here, *t* is the time elapsed from the entry of the TCR to the close contact and 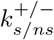 are the on/off rates for self/non-self ligands binding to TCR.

The initial conditions for this system of PDEs for are

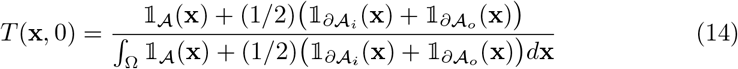

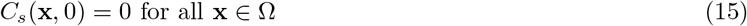

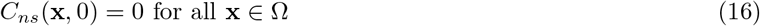

where 𝟙_𝒮_ (**x**) is the indicator function of the specified set 𝒮. The set 𝒜 is defined as

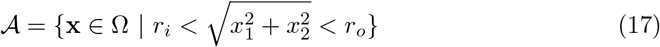

with inner radius *r*_*i*_ and outer radius *r*_*o*_ such that *r*_*i*_ *< r*_*o*_ *< R*. The sets *∂*𝒜_*i*_ and *∂*𝒜_*o*_are the inner and outer boundaries of 𝒜 defined as

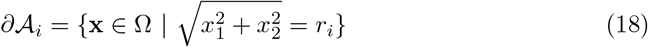

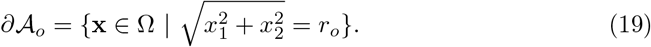

The boundary conditions are

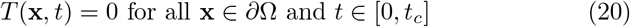

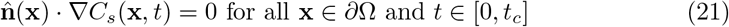

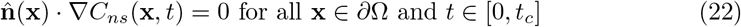

where 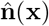 is the outward-facing unit normal vector at **x** *∈∂*Ω. These boundary conditions represent the ability of unbound TCRs to leave the close contact and the inability of bound TCRs to leave the close contact.

We used the py-pde Python package [30] to numerically solve the system of PDEs in Eqs. 11-13 using finite difference methods.

### TCR activation model

From a set of replicate trajectories generated from stochastic simulations, we can calculate the probability that over the lifetime of the close contact at least one TCR will reach the fully phosphorylated state. While a TCR is in the close contact, the functional TCR complex can assemble and be phosphorylated. In our model, we ignore complex assembly and only focus on phosphorylation. If the phosphatase concentration is low enough in the close contact, we can assume that phosphorylation occurs through *n* irreversible steps. Then, the probability that a single TCR complex is fully phosphorylated by time *t* in *n* irreversible steps, each with the same rate *k*_*p*_, follows the Erlang distribution

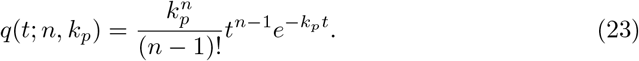

If we were to allow for dephosphorylation, we could write the model as a linear system of ordinary differential equations where the fully phosphoryalted state is absorbing.

However, in this work we assume that dephosphorylation in the close contact occurs at a slow enough rate to be ignored. In Fig 3, we also calculate activation probabilities for

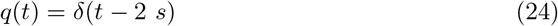

as was used in Fernandes *et al*. [22].

In order to calculate the probability of a single TCR complex reaching the fully phosphorylated state before leaving the close contact for KS-KP, given the fraction of ligands that are non-self, we calculate

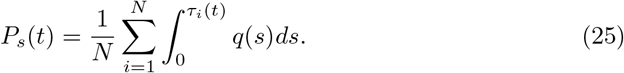

with

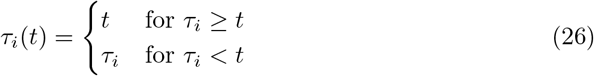

where *τ*_*i*_ is the close contact residence time of the TCR in trajectory *i*. Eq. 25 is the probability of a single TCR complex reaching the fully phosphorylated state before leaving the close contact, averaged over all trajectories. We calculated the standard deviation of *P*_*s*_(*t*) for KS-KP according to

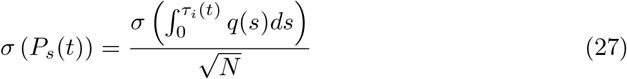

given the independence of *τ*_*i*_ between different trajectories.

To calculate the probability of a TCR complex reaching the fully phosphorylated state with cKP, we use the same trajectories but instead calculate

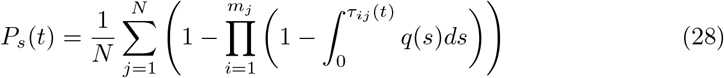

where the product is over *m*_*j*_ bound periods, indexed by *i*, for a single trajectory indexed by *j*. Each term of the product is the probability that in trajectory *j* during the *i*^th^ bound period the TCR did not reach the fully phosphorylated state. The length of the *i*^th^ bound period of trajectory *j* is

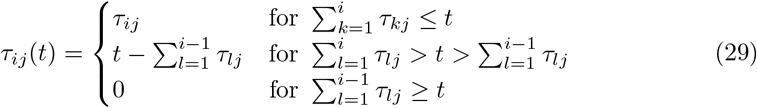

which allows us to consider arbitrary times, including those which fall in the middle of bound period for some trajectory. The product is the probability that none of the bound periods in trajectory *j* resulted in the TCR reaching the fully phosphorylated state, so that one minus this quantity is the probability that at least one of the bound periods resulted in full phosphorylation. This quantity is then averaged over all *N* trajectories. As with *P*_*s*_(*t*) for KS-KP, we calculated the standard deviation of *P*_*s*_(*t*) for cKP according to

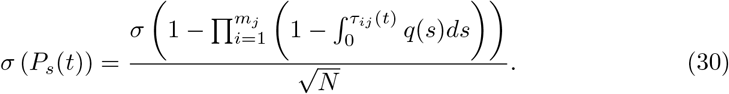

Eqs. 25 & 28 provide the probability of a *single* TCR complex reaching the fully phosphorylated state by time *t*. However, in an actual T cell multiple TCRs can enter and leave a close contact, and we are interested in the probability that *any one* of those TCRs become fully phosphorylated in the lifetime of the close contact. We can write the probability that no TCR will enter the close contact and become fully phosphorylated over the close contact lifetime by assuming a constant rate of TCR entry into the close contact and by considering *m* – 1 time intervals of length *t*_*c*_*/m*, for positive integer *m*. The entry rate of TCR entry into the close contact is [22]

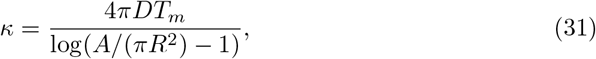

where *T*_*m*_ is the bulk membrane TCR density far from the close contact, *A* is the cell surface area, and *R* is the radius of the close contact. Using Eq. 31 and discretizing time into intervals of length *t*_*c*_*/m*, the probability that a TCR will not activate is

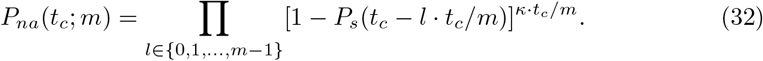

Taking the limit of *P*_*na*_(*t*_*c*_; *m*) as *m → ∞*, corresponding to continuous time, we have the geometric integral [31]

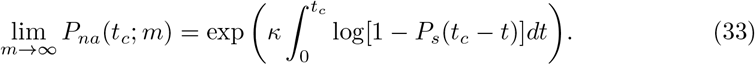

Because Eq. 33 provides the probability that no TCR will activate in *t*_*c*_, we can then easily find the probability that *at least one* TCR will activate in *t*_*c*_ as

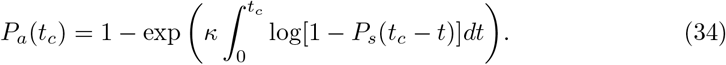

This is the activation probability shown in Figs. 3-5. While the notation of our model obscures the dependence of *P*_*a*_(*t*_*c*_) on *G*, we emphasize that with a fixed value of *M, G* fully determines the self and non-self concentrations *M*_*s*_ and *M*_*ns*_ and through the stochastic simulations influences *P*_*a*_(*t*_*c*_). In order to find the standard deviation of *P*_*a*_(*t*_*c*_), we used linear error propagation theory implemented in the Uncertainties Python package [32] using Eqs. 30 & 27 for the standard deviation of *P*_*s*_(*t*) for cKP and KS-KP, respectively.

## Acknowledgments

A.S.M. and A.W.E. were partially supported by the United States Defense Advanced Research Projects Agency (http://www.darpa.mil/) RadioBio program under grant number HR001117C0125, and by a Discovery grant from the Natural Sciences and Engineering Research Council of Canada (http://www.nserc.ca/). K.A.G. acknowledges support by the WISE program of the Dutch Research Council (NWO).

The funders had no role in study design, data collection and analysis, decision to publish, or preparation of the manuscript.

