## Supplemental information for "Comparing kinetic proofreading and kinetic segregation for T cell receptor activation"

Table S1: Model parameters used in main text figures.

| Parameter (Unit) | Fig. 1 | Fig. 2 | Fig. 3 | Fig. 4 | Fig. 5 | Fig. 6 | Fig. 7 |
|---|---|---|---|---|---|---|---|
| $\Delta t$ ($s$) | $10^{-4}$ | $10^{-4}$ | $10^{-4}$ | $10^{-4}$ | $10^{-4}$ | $10^{-4}$ | $10^{-4}$ |
| $D$ ($\mu m^2 \cdot s^{-1}$) | 0.052 | 0.052 | 0.052 | 0.052 | 0.052 | 0.052 | 0.052 |
| $M$ ($\mu m^{-2}$) | 330 | 330 | 330 | 330 | 330 | 330 | 330 |
| $G$ (Unitless) | 0.1 | 0-0.1 | 0-0.1 | 0, 0.01, 0.04 | 0-0.1 | 0, 0.01, 0.1 | 0, 1 |
| $R$ ($\mu m$) | 0.22 | 0.22 | 0.22 | 0.22 | 0.22 | 0.22 | 0.22 |
| $T_m$ ($\mu m^{-2}$) | - | 100 | 100 | 100 | 100 | 100 | 100 |
| $A$ ($\mu m^2$) | - | 415 | 415 | 415 | 415 | 415 | 415 |
| $t_c$ ($s$) | - | 60 | 5, 30, 120 | 0-120 | 5, 30, 120 | 1-120 | 60 |
| $k_s^+$ ($\mu m^2 \cdot s^{-1}$) | 0.1 | 0.1 | 0.1 | 0.1 | 0.1 | 0.1 | See Table S2 |
| $k_s^-$ ($s^{-1}$) | 50 | 50 | 50 | 50 | 50 | 50 | See Table S2 |
| $k_{ns}^+$ ($\mu m^2 \cdot s^{-1}$) | 0.1 | 0.1 | 0.1 | 0.1 | 0.1 | 0.1 | See Table S2 |
| $k_{ns}^-$ ($s^{-1}$) | 1 | 1 | 1 | 1 | 0.1, 1, 10 | 0.1, 1, 10 | See Table S2 |
| $k_p$ ($s^{-1}$) | 1 | 1 | 1 | 1 | 1 | 1 | 1 |
| $n$ (Unitless) | 3 | 2, 3, 4 | 3 | 3 | 3 | 3 | 3 |

Table S2: Ligands used in Fig. 7.

| Ligand name | Ligand type | $k_{s/ns}^+$ ($\mu m^2 \cdot s^{-1}$) | $k_{s/ns}^-$ ($s^{-1}$) | $EC_{40}$ or $EC_{50}$ ($\mu M$) | Reference |
|---|---|---|---|---|---|
| OVA | Non-self | 1.2 | 10.8 | $EC_{50}$: $3.33 \cdot 10^{-5}$ | Huang *et al.* (2010) [25] |
| A2 | Non-self | $3.5 \cdot 10^{-1}$ | 4.7 | $EC_{50}$: $6.26 \cdot 10^{-4}$ | Huang *et al.* (2010) [25] |
| Hb | Non-self | $2.3 \cdot 10^{-1}$ | 1.22 | $EC_{40}$: $4 \cdot 10^{-4}$ | Hong *et al.* (2015) [26] |
| V-OVA | Self | $4 \cdot 10^{-4}$ | 1.4 | $EC_{50}$: 9.09 | Huang *et al.* (2010) [25] |
| R4 | Self | $4.4 \cdot 10^{-4}$ | 2.6 | $EC_{50}$: 4 | Huang *et al.* (2010) [25] |
| I72 | Self | $1.45 \cdot 10^{-2}$ | 2.99 | $EC_{40}$: 7 | Hong *et al.* (2015) [26] |
| A72 | Self | $1.29 \cdot 10^{-2}$ | 3.79 | $EC_{40}$: 60 | Hong *et al.* (2015) [26] |

Table S3: PDE model parameters used in Figure S2.

| Parameter (Unit) | Value |
|---|---|
| $\Delta t$ ($s$) | $10^{-4}$ |
| $D$ ($\mu m^2 \cdot s^{-1}$) | 0.052 |
| $M$ ($\mu m^{-2}$) | 330 |
| $R$ ($\mu m$) | 0.22 |
| $r_i$ ($\mu m$) | 0.21 |
| $r_o$ ($\mu m$) | 0.2125 |
| Number of $G$ values | 21 |
| PDE solution step size | $10^{-5}$ |
| PDE solution time interval | 0.125 |
| Number of PDE grid points | 140 |

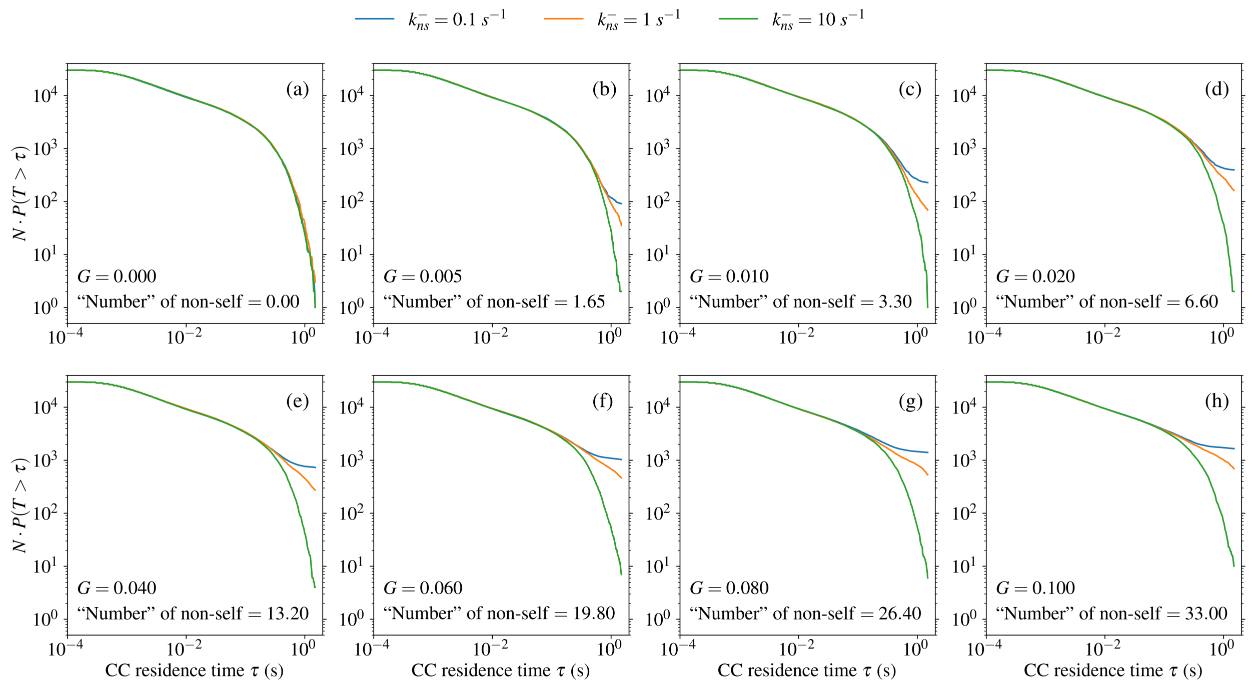


Figure S1: Single TCR close contact (CC) lifetime distributions from simulations for different values of $G$ and non-self off rate, $k_{ns}^-$. The y-axes are the total fraction of simulations for each $k_{ns}^-$ at a given value of $G$ that stayed in the close contact for more than $\tau$ seconds, equivalent to the total number of simulations for each $k_{ns}^-$ and $G$ times the empirical probability $P(T > \tau)$. In (a), there is no non-self ligand ($G = 0$), so the three $k_{ns}^-$ values yield nearly identical $N \cdot P(T > \tau)$ curves as expected (as $N \to \infty$, the three curves should converge toward the exact same curve). When $G = 0.005$ (b), where the approximate number of ligands implied ($G \cdot M$) is 1.65, there is a clear difference in the number of trajectories that stay in the close contact for over 1 second. Subfigures (c)-(h) demonstrate the increasing probability of long close contact lifetimes as $G$ increases, as well as the relatively constant fold differences in long close contact residence time probabilities between the three $k_{ns}^-$ values. There is a striking consistency in $N \cdot P(T > \tau)$ for $\tau < 1$ s across all $k_{ns}^-$ and $G$ values, indicating that ligand binding plays little role in determining short TCR close contact residence times. However, the different $N \cdot P(T > \tau)$ curves for different $k_{ns}^-$ values do diverge at shorter $\tau$ values as $G$ increases, due to larger total binding rates ($G \cdot M \cdot k_{ns}^+$) which mean that non-self ligands can influence earlier events in TCR diffusion.

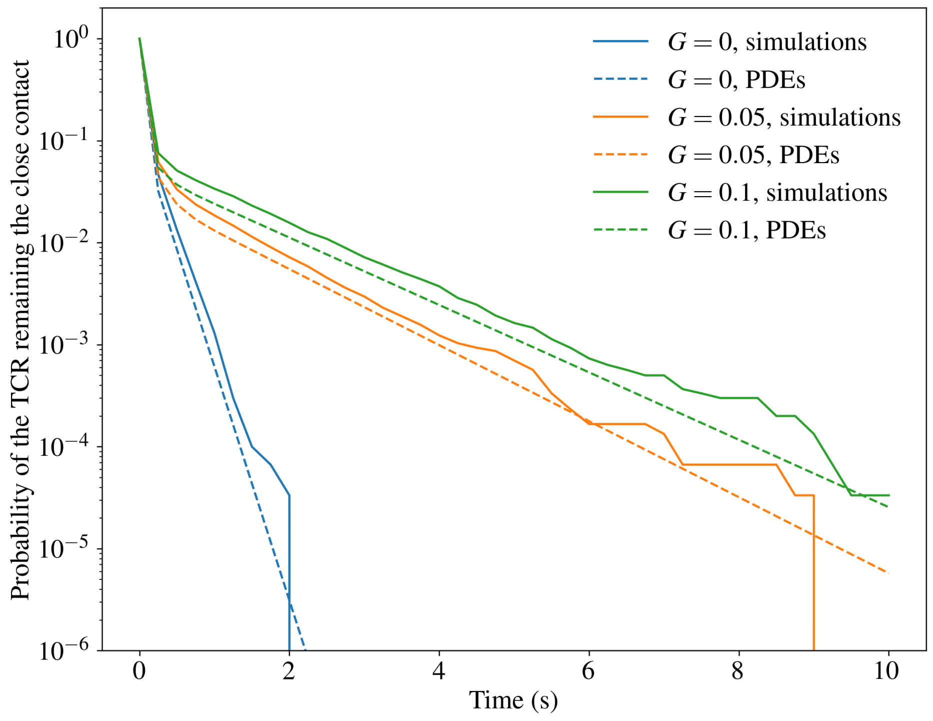


Figure S2: Consistency check between our model (labelled *simulations*) and the PDE-based model from Fernandes *et al.* [22] (labelled *PDEs*), demonstrating the consistency of the two models in predicting the probability of the TCR remaining in the close contact with respect to time.

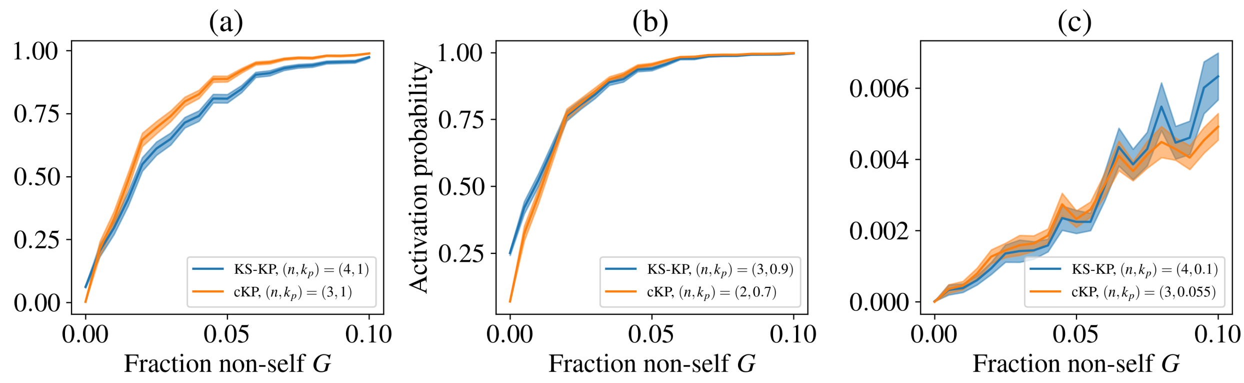


Figure S3: Similar solutions for activation probability as a function of $G$ can be found for both KS-KP and cKP: since a similar curve can be produced for both KS-KP and cKP for different parameters, it is difficult to distinguish between the models if only activation probability is considered. Subfigure (a) is an expanded version of the inset in Figure 2, with $(n, k_p) = (4, 1)$ for KS-KP and $(n, k_p) = (3, 1)$ for cKP. The other subfigures show even better fits: in subfigure (b) we have $(n, k_p) = (3, 0.9)$ for KS-KP and $(n, k_p) = (2, 0.7)$ for cKP, while in subfigure (c) we have $(n, k_p) = (4, 0.1)$ for KS-KP and $(n, k_p) = (3, 0.055)$ for cKP. Similar solutions can be found throughout the parameter space, and these particular values are provided for illustration.

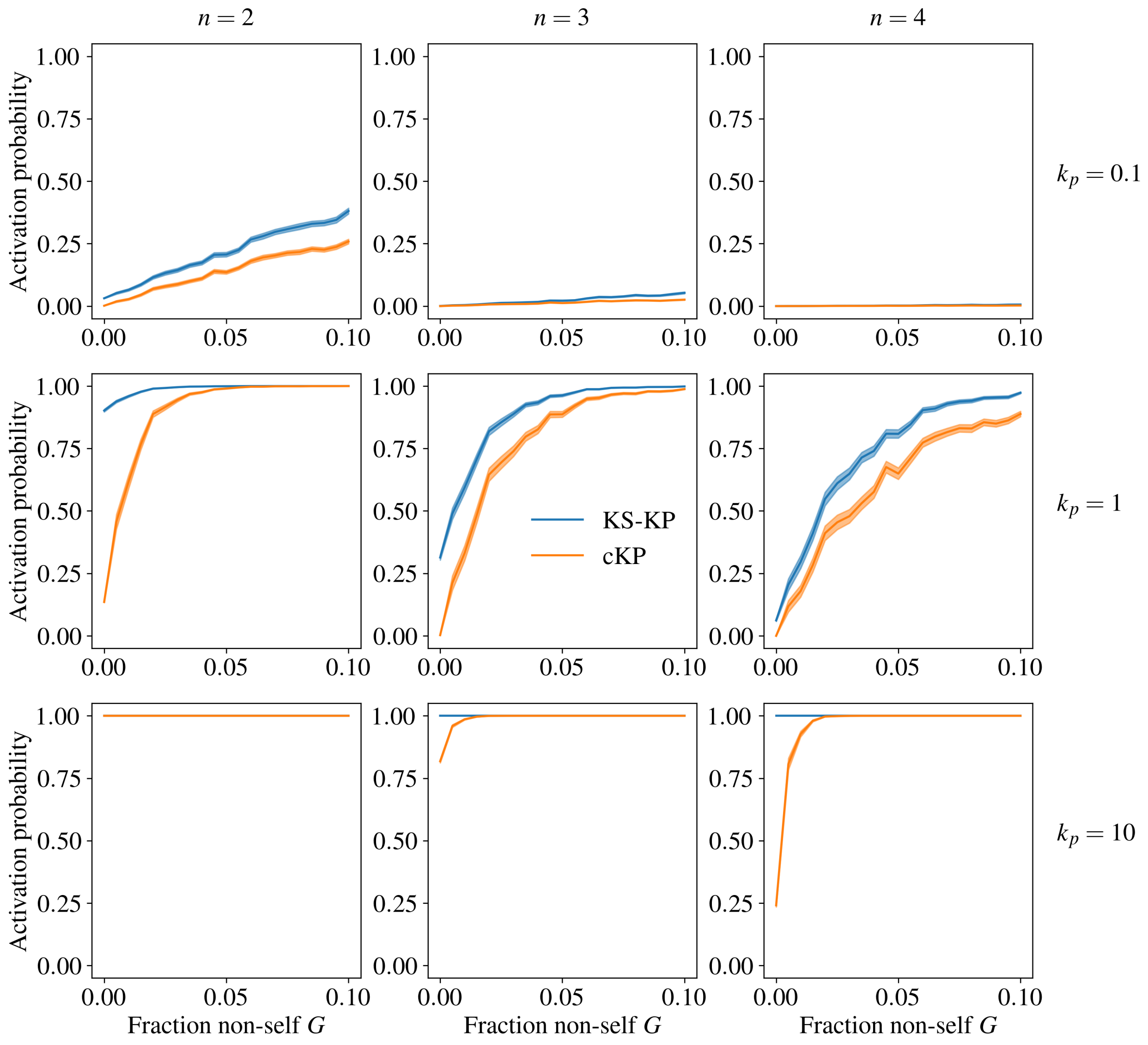


Figure S4: An expanded version of Fig. 2, plotting activation probability against fraction of non-self ligands $G$. Compared with Fig. 2, the effect of different phosphorylation rates $k_p$ is illustrated: at low $k_p$, the T cell rarely activates, while at high $k_p$, it almost always activates (in both cases, somewhat dependent on the number of phosphorylation steps $n$). To allow direct comparison, the middle row ($k_p = 1$) is copied from Fig. 2.

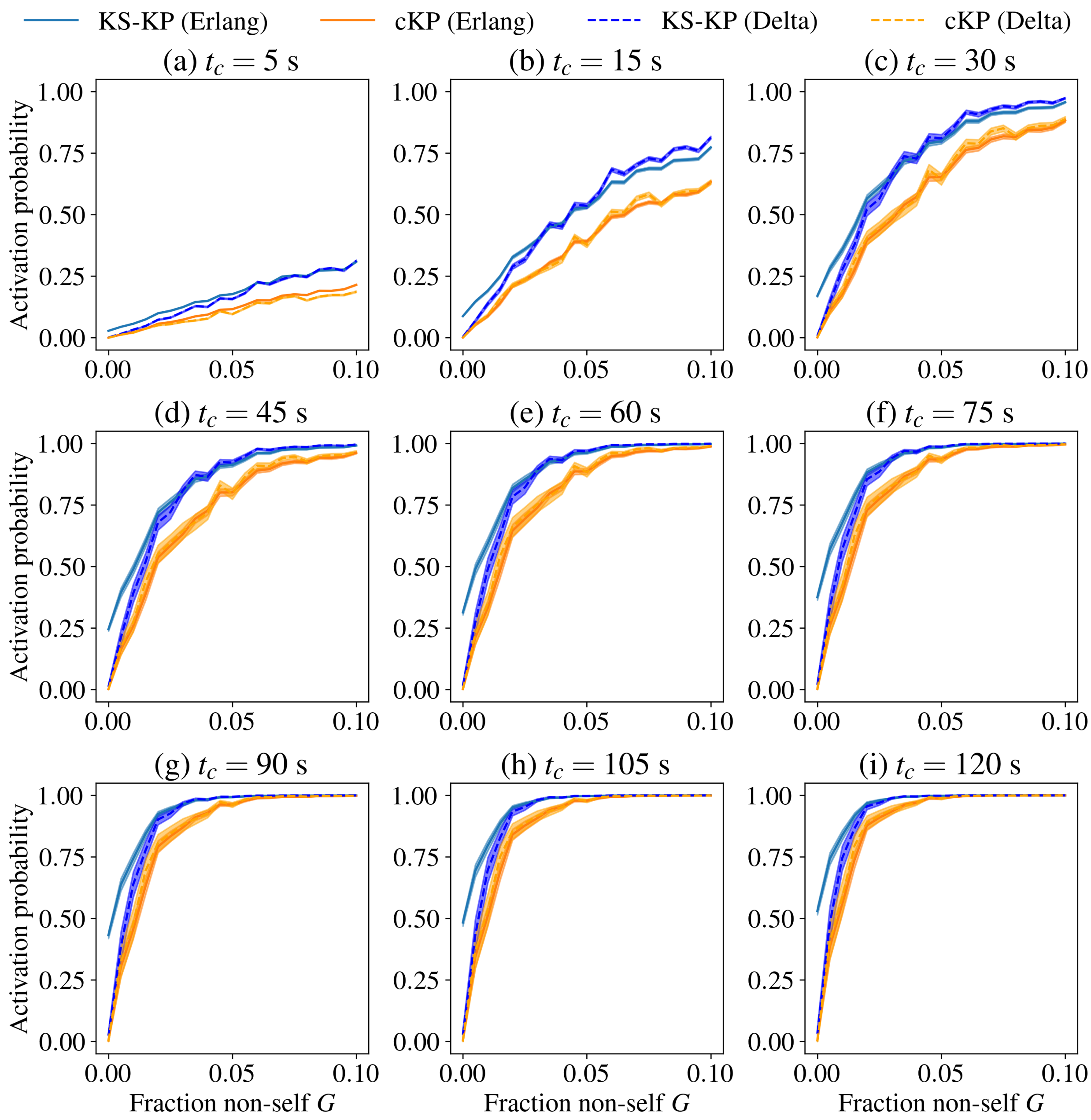


Figure S5: An expanded version of Fig. 3, plotting activation probability against fraction of non-self ligands $G$ under the Erlang and Delta activation models. A steady increase in activation probability is observed as $t_c$ increases. It should be noted that only the KS-KP Erlang model has a significant activation probability as $G \to 0$, regardless of $G$. To allow direct comparison, the plots for $t_c = 5$ s, 30 s, and 120 s, in subfigures (a), (c), and (i) respectively, are copied from Fig. 3.

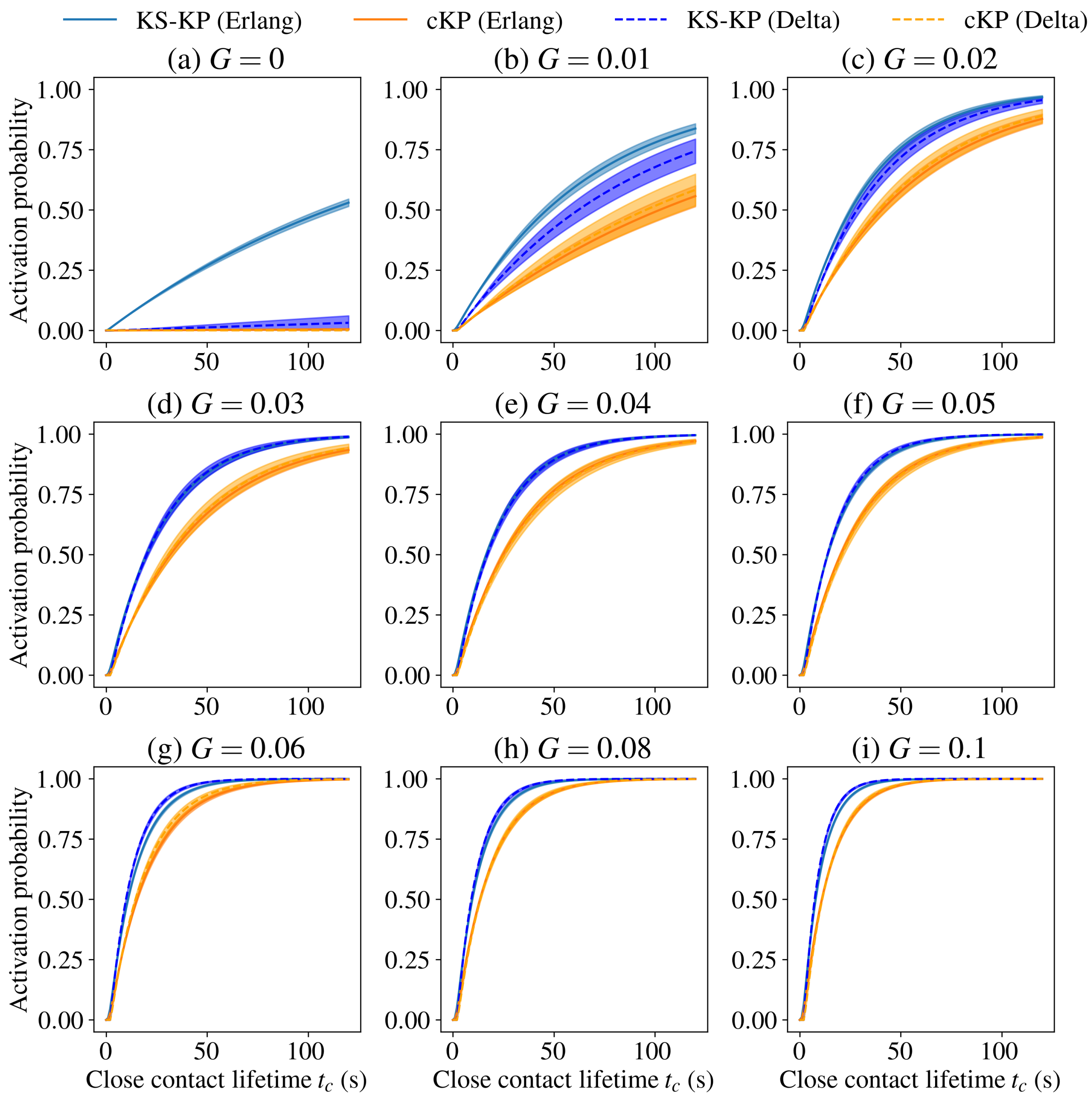


Figure S6: An expanded version of Fig. 4, plotting activation probability against close contact lifetime $t_c$ under the Erlang and Delta activation models. Large differences are seen between KS-KP (Erlang) and the other models at low $G$, though the difference rapidly diminishes with increasing $G$. To allow direct comparison, the plots for $G = 0$, 0.01, and 0.04, in subfigures (a), (b), and (e) respectively, are copied from Fig. 4.

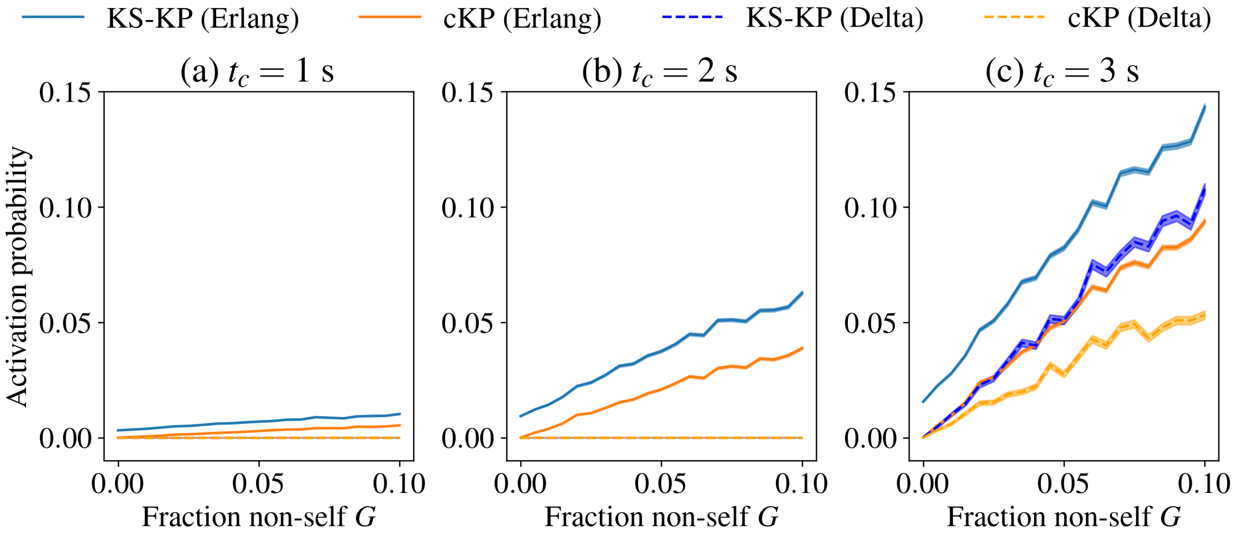


Figure S7: For $t_c \leq 2$ s, the Erlang model indicates a significant probability of activation (up to $p = 0.06$, subfigure (b)). Meanwhile, since the Delta model predicts immediate, full activation after a ligand is present in the close contact for more than 2 s, the probability of activation in this model is zero for $t_c \leq 2$ s (subfigures (a), (b)). A gap persists for all $G$ at $t_c = 3$ s, subfigure (c), but rapidly shrinks as $t_c$ increases (Fig. S5).

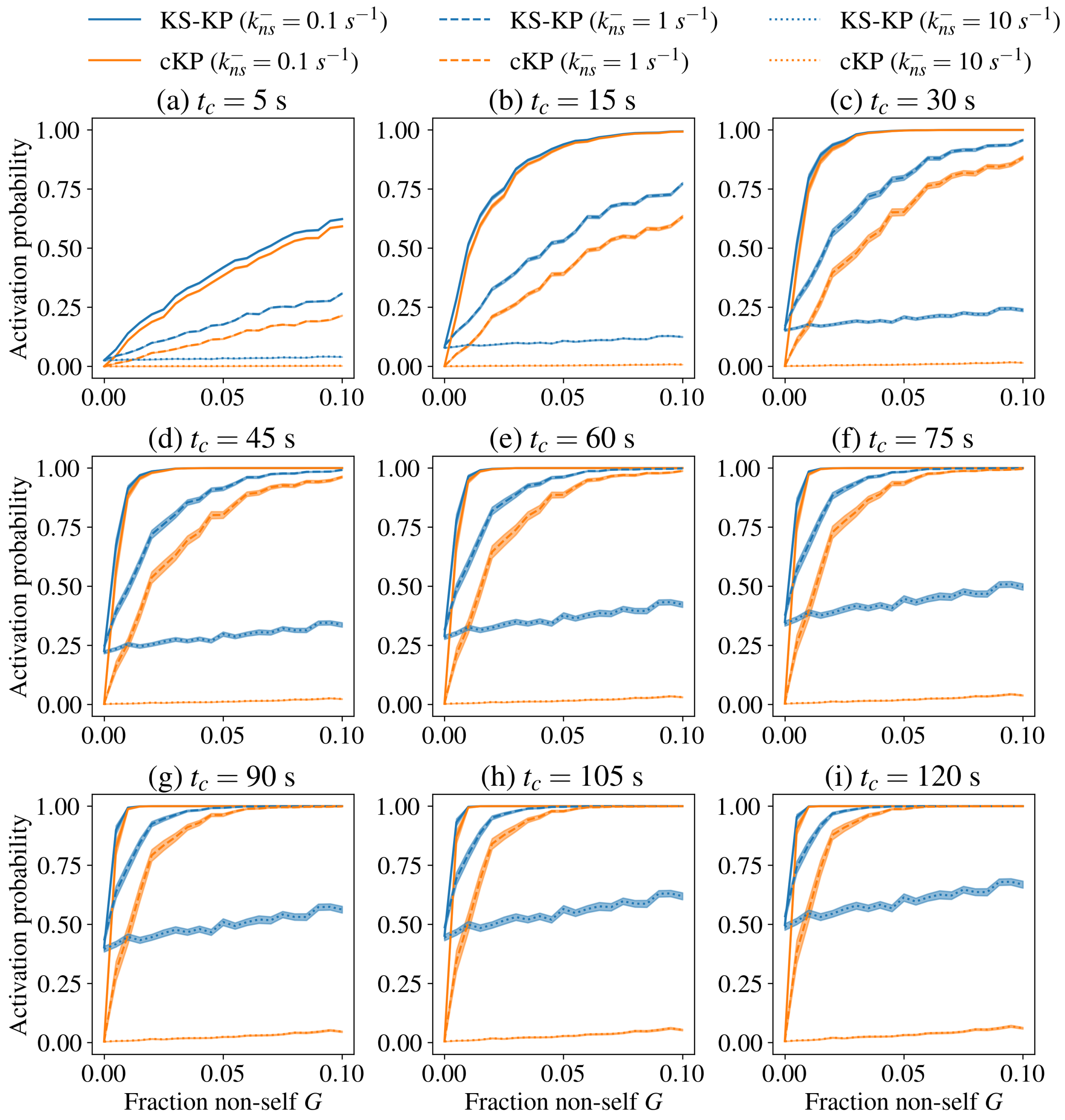


Figure S8: An expanded version of Fig. 5, plotting activation probability against fraction non-self $G$ for various non-self off rates $k_{ns}^{-}$ and close contact lifetimes $t_c$. Large differences are seen between KS-KP and cKP at low $G$, and the differences persist as $G$ increases for large values of $k_{ns}^{-}$. To allow direct comparison, the plots for $t_c = 5$ s, 30 s, and 120 s, in subfigures (a), (c), and (i) respectively, are copied from Fig. 5.

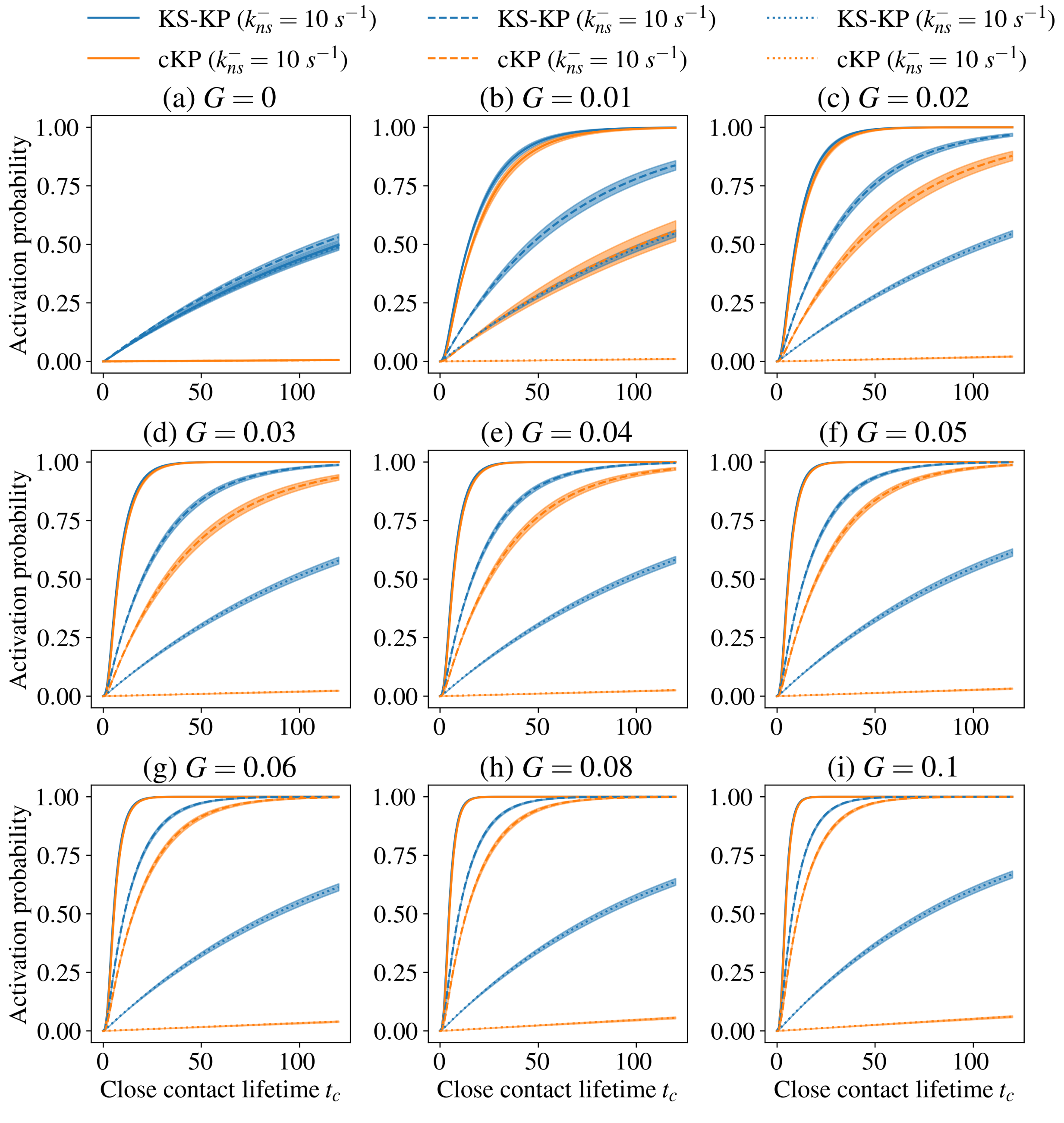


Figure S9: An expanded version of Fig. 6, plotting activation probability against close contact lifetime $t_c$ for various non-self off rates $k_{ns}^{-}$ and non-self fractions $G$. Similarly to Fig. S8, large differences are seen between KS-KP and cKP at low $G$. To allow direct comparison, the plots for $G = 0$, 0.01, and 0.1, in subfigures (a), (c), and (i) respectively, are copied from Fig. 6.
